# dagLogo: an R/Bioconductor Package for Identifying and Visualizing Differential Amino Acid Group Usage in Proteomics data

**DOI:** 10.1101/2020.08.28.254623

**Authors:** Jianhong Ou, Haibo Liu, Niraj K. Nirala, Alexey Stukalov, Usha Acharya, Michael R. Green, Lihua Julie Zhu

## Abstract

Sequence logos have been widely used as graphical representations of conserved nucleic acid and protein motifs. Due to the complexity of the amino acid (AA) alphabet, rich post-translational modification, and diverse subcellular localization of proteins, few versatile tools are available for effective identification and visualization of protein motifs. In addition, various reduced AA alphabets based on physicochemical, structural, or functional properties have been valuable in the study of protein alignment, folding, structure prediction, and evolution. However, there is lack of tools for applying reduced AA alphabets to the identification and visualization of statistically significant motifs. To fill this gap, we developed an R/Bioconductor package dagLogo, which has several advantages over existing tools. First, dagLogo allows various formats for input sets and provides comprehensive options to build optimal background models. It implements different reduced AA alphabets to group AAs of similar properties. Furthermore, dagLogo provides statistical and visual solutions for differential AA (or AA group) usage analysis of both large and small data sets. Case studies showed that dagLogo can better identify and visualize conserved protein sequence patterns from different types of inputs and can potentially reveal the biological patterns that could be missed by other logo generators.

## Introduction

Since their first introduction in 1990, sequence logos have been widely used to visualize conserved patterns among large sets of nucleic acid and peptide sequences [1–4]. Dozens of sequence logo generators with different functionalities and performances have been developed; these generators require various types of inputs and background models, and use different algorithms and graphical representations [1–3, 5–15]. A comprehensive list of these tools, along with their features, is available in **Supplementary Table S1**. Almost all tools provide no alignment utilities but require multiple sequence alignments in various formats, or position weight/frequency matrices as inputs. A majority of these tools are also based on information theory without statistical significance tests [16–18]. More than two thirds of the existing tools cannot effectively display both over- and under-represented residues (**Supplementary Table S1**), even though the information on under-represented residues may be as valuable as that of over-represented residues [12, 19].

It is known that both background models and sizes of input sets affect the identification and visualization of sequence patterns [3, 7, 18, 20]. However, a majority of logo generators do not allow users to choose a dataset-specific background model. For those that do allow it, the only option for a majority of them is a background model based on global residue frequencies or GC%, which can be specified by users, calculated from input sequences, or derived from a species-specific whole genome or proteome (**Supplementary Table S1**). These simple background models usually work well for identifying and visualizing nucleic acid patterns due to the simple nucleotide alphabet and limited modifications of nucleic acids, especially DNA. However, in many cases, they may not be optimal for protein pattern discovery because of the complexity of the amino acid (AA) alphabet, rich post-translational modification of proteins, diverse subcellular localization, cell or tissue type-specific expression profiles, and experimental protocol-specific biases. To our knowledge, only Two Sample Logo [13], iceLogo [7, 21], pLogo [6], kpLogo [22], and PTM-Logo [23] offer some advanced options of background models. Among them, iceLogo offers the most choices, i.e., i) species-specific, proteome-wide AA frequencies as the background; ii) position-specific AA frequencies as the background; and iii) an experimental protocol bias-aware background.

To date, there are no tools that have the functionalities to group AAs based on their physicochemical or other properties and test the differential usage of AA groups. Historically, twenty naturally occurring AAs has been classified into two to 19 groups based on some measures of their relative similarity in physicochemical, structural, and functional properties; this has resulted in a variety of reduced (degenerate or simplified) alphabets [24–27]. With reduced alphabets, several AAs of a similar property are lumped together and represented by one letter. Reduced AA alphabets have been shown to be valuable for protein pattern discovery [28–30] and in the study of protein alignment, folding, structure prediction, and evolution [25, 31–38]. For example, serine (Ser) and threonine (Thr) are highly interchangeable as phosphorylation sites of Ser/Thr kinase substrates [39], and thus the two AAs can be binned into the same degenerate group to reduce the complexity of resulting logos showing conserved patterns of Ser/Thr kinase (STK) substrates. Thus, unsurprisingly, coloring AA residues based on their similarity of physicochemical properties has been a general practice of many logo generators long before [2] (**Supplementary Table S1**). However, applying reduced AA alphabets to motif identification and visualization had rarely been practiced until recent [28, 29]. This limitation is more obvious for sequence patterns of proteins than sequence patterns of nucleic acids because protein sequences have a greater alphabet complexity and variability of the occurrences of individual residues. Even though RaacLogo [28] and logoJS [29] allow degenerate representations of reduced AA alphabets [40, 41], both were based on information theory without statistical significance tests, and most importantly, neither offers alternative background models (**Supplementary Table S1**).

Taken together, none of the existing tools has implemented all the following functionalities: allowing flexible formats for input sets, providing comprehensive options to build optimal background models, and implementing reduced AA alphabets to group AAs of similar properties. To fill this gap, we have developed an open source R/Bioconductor package dagLogo for identifying and visualizing conserved peptide patterns with the probability theory. In addition to having the functionality to generate various background models, dagLogo also accepts aligned subsequences or unaligned subsequences of different lengths as input, and it visualizes significantly over- and under-represented AA residues or AA residue groups in experimental context-aware ways. We demonstrate here that dagLogo can better identify and visualize conserved patterns hidden in peptide sequences regardless of length and alignment status, and can potentially reveal patterns that could be missed by other logo generators.

## Results

### Case study 1: dagLogo successfully identifies the substrate sequence specificity of granzyme B at both individual AA and physicochemical property levels

Granzyme B (GRB) is an apoptotic serine protease found in secretory granules of cytotoxic T lymphocytes and natural killer cells [42, 43]. Early *in vitro* screening studies revealed that GRB preferentially cleaves the peptide bonds immediately proceeded by aspartate residues at P1 positions [44], and determinants of its substrate specificity span from residues at P4 to P2’ (for notation Px and Px’, see **Figure 1**) [45]. Harris *et al*. identified an optimal consensus substrate of GRB with (I/V)-(E/Q/M)-X-(D^↓^X)-G at positions P4 to P2’, where X is any AA residue and the peptide bond between D and X is the GRB cleavage site [45]. Later, an *in vivo* study identified the proteome-wide substrates for the human as well as the mouse GRB [46], which not only confirmed the *in vitro* findings about the determinants of GRB substrate specificity but also revealed that more residues from P3’ to P9’ are related to substrate specificities and cutting efficiency [46]. Van Damme *et al*. further revealed the physicochemical property constraints in the substrate sequence pattern in terms of charge, hydrophobicity, and side chain size of residues [46]. GRB prefers negatively charged AAs at P3 and P2’-P8’, and slightly dislikes positively charged AAs around the cleavage site. They also found that AAs at P4 tend to be hydrophobic, followed by less hydrophobic AA at P3. Additionally, AAs at positions P3, P2, and P1’ flanking the cleavage site tend to be small residues [46]. These observations are consistent with the molecular structure of the GRB substrate binding pocket [47–49].

**Figure 1.**
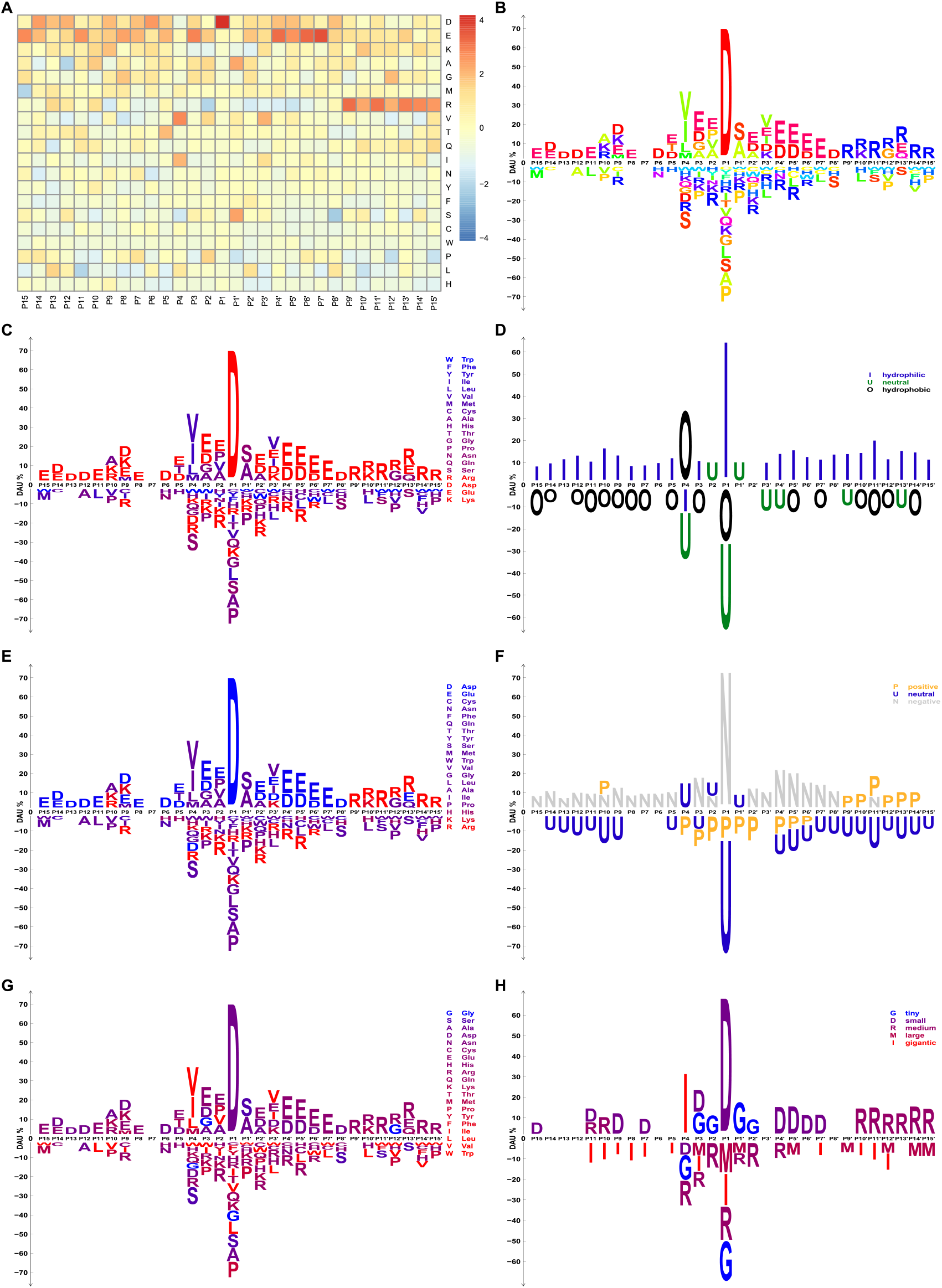
Heatmap and sequence logos showing the substrate sequence pattern of the human granzyme B. A background model for *Z*-test was built from randomly sampled subsequences of 30 AA residues from the UniProt human reference proteome. The significance level is set as 0.05 for DAU tests. (**A**) *Z*-scores of DAU tests for each residue at each pattern position are shown in the heatmap. (**B, C, E,** and **G**) Over- and under-represented AA residues with significant DAU at each pattern position are shown as sequence logos. Individual AA residues are colored differently and according to their hydrophobicity indices, pI indices, or sizes, respectively. (**D, F,** and **H**) AA residues are grouped and collapsed based on their hydrophobicity indices, charge status under physiological conditions, and sizes, respectively. Over- and under-represented AA residue groups with significant DAU at each pattern position are shown as degenerate logos. AA residue groups are colored according to their hydrophobicity indices: (**D**) hydrophobic AAs = {W, F, Y, L, I, V, M, C}, neutral AAs = {H, A, T, P, G, N, A, S}, and hydrophilic AAs = {R, K, D, E}; their charge status under physiological conditions: (**F**) positively charged AAs = {H, K, R}, neutral AAs = {A, C, F, G, I, L, M, N, P, Q, S, T, V, W, Y}, and negatively charged AAs = {D, E}; or sizes: (**H**) tiny = {G, S, A}, small = {D, N, C, E, H}, medium = {R, Q, K, T}, large = {M, P, Y, F}, and gigantic = {I, L, V, W}. Van Damme *et al.* thinks that the weak arginine conservation from P11’ to P15’ is probably due to technical artifacts caused by the *N*-terminal COFRADIC sorting procedure [46].

We reanalyzed the proteome-wide substrate data of the human GRB [46] using our dagLogo package and compared the resulting logo with logos generated by other common tools with the same input and background sets of subsequences, wherever possible. **Supplementary Figure S1** displays the sequence patterns among the human GRB substrates revealed by dagLogo, iceLogo, pLogo, and WebLogo. Although all logo generators successfully identified the major determinant of substrate specificity, i.e., enriched Asp in P1, both pLogo and WebLogo failed to reveal the important determinants flanking P1 (**Supplementary Figures S1C** and **S1D**). pLogo calculates the log odds, log (*p*/(1 – *p*)), where *p* is the binomial probability of an AA residue occurring at a given position. pLogo emphasizes the big differences but downplays the minor differences. This may explain why residues flanking P1, which contributes less to substrate specificity, are not as clearly represented. WebLogo does not allow users to provide a background model. As a result, a uniform background frequency of 20 AA residues was used as the background model by default. In addition, it can only display positively scored residues based on the information theory. Logos generated by dagLogo (**Supplementary Figure S1A**) and iceLogo (**Supplementary Figure S1B**) are very similar, which clearly displays both the major and minor determinants of the substrate specificity. The subtle differences might be because the two tools calculate the variances differently (see **Discussion**).

Next, we performed more analyses of the human GRB substrate specificity, using dagLogo to display other important features of our package. **Figure 1A** displays the differential usage of each AA residue from P15 to P15’ through a heatmap. This heatmap gives a detailed view of AA distribution at each position in comparison with the given background model. **Figures 1B, 1C, 1E, and 1G** display significant differential usages of individual AAs at each position, where AA residues are colored individually or collectively according to their hydrophobicity indices, pI indices, and sizes, respectively. By contrast, **Figures 1D, 1F, and 1H** show significant differential usages of AA groups using degenerate logos, where AAs sharing a similar physicochemical property are grouped using the reduced AA alphabets and are tested for differential usage. dagLogo-based analyses, as displayed in **Figure 1,** recapitulated the previous *in vitro* and *in vivo* findings. These findings not only confirm the properties of GRB substrate specificity in terms of sequence preferences at individual AA level, but also illuminate the restrictions in physical properties like hydrophobicity, charge, and sizes of residues [44–46].

### Case study 2: dagLogo successfully differentiates substrate specificities of three human Ser/Thr protein kinases

One of the most common, important, and dynamically reversible post-translation modifications is phosphorylation, which is controlled by various protein kinases and phosphatases [50–52]. Protein kinases employ combinatorial mechanisms to define their substrate specificities, including substrate peptide specificity and substrate recruitment specificity [53, 54]. Substrate motifs of protein kinases have been identified over the past 70 years, though different studies have resulted in more or less different forms of motifs [55–59] (https://www.phosphosite.org). According to the AA residues phosphorylated by protein kinases, eukaryotic kinases are classified into Ser/Thr kinases (STKs), Tyr kinases, and dual-specificity kinases, which can phosphorylate Ser, Thr, and Tyr residues. At least 125 of the 568 human protein kinases are STKs [60, 61], and more than 86% of phosphorylated residues are Ser, followed by 11.8% being Thr and 1.8% being Tyr in the human HeLa cells [62].

PKACA (cAMP-dependent protein kinase catalytic subunit alpha) and PKCA (protein kinase C alpha) are members of the AGC protein kinase superfamily. The consensus substrate motif of PKACA is (R/K)(R/K)X(S*/T*), where S*/T* is the phosphorylated Ser or Thr [63, 64] (https://www.phosphosite.org). It prefers hydrophobic AAs at position +1 and basic AAs at positions −7 to −2. PKCA, like other PKC family members, preferentially phosphorylate peptides with hydrophobic AAs at position +1. It also favors substrates with basic residues at positions −7 to −1, and +2 to +4 [65]. An established consensus substrate motif for PKCA is (R/K)(R/K)X(S/T)X(R/K) [65].

In contrast to most STKs whose substrate phosphorylation sites are preferentially flanked by basic AA residues, sequences flanking the canonical substrate phosphorylation sites of the protein kinase CK2A1 (catalytic subunit of a constitutively active serine/threonine-protein kinase, Casein Kinase 2 (CK2)) are preferentially occupied by acidic residues [66, 67]. A consensus CK2 substrate motif is (S*/T*)(D/E)X(E/D) [67–69]. Acidic residues at positions +1 and +3 are the most important specificity determinants, but additional acidic residues at positions from −2 to +7 and even further contribute to substrate specificity too. Basic and bulky hydrophobic residues at these positions are depleted [66, 67]. Besides local sequence determinants, the overall protein conformation and accessibility of candidate substrate can affect phosphorylation efficiency of the substrates by CK2 [66]. Additionally, CK2 can use phosphoserine residues as consensus specificity determinants to phosphorylate non-canonical substrates via hierarchical phosphorylation [70].

Using a background set which consisted of randomly sampled peptide sequences of 15 AA residues from the human proteome we reanalyzed the substrate phosphorylation motifs of three human STKs—PKACA, PKCA, and CK2A1—using dagLogo. The substrate sequence patterns identified for PKACA (**Figure 2A**), PKCA (**Figure 2B**), and CK2A1 (**Figure 2C**) by dagLogo are largely consistent with those curated by the PhosphoSitePlus database (https://www.phosphosite.org). Interestingly, we detected significant frequencies of serine residues flanking the phosphorylation sites of CK2A1, where acidic residues are usually preferred. This is consistent with the hierarchical phosphorylation property of CK2 [70]. The substrate motif of PKACA is much more similar to that of PKCA than to that of CK2A1.

**Figure 2.**
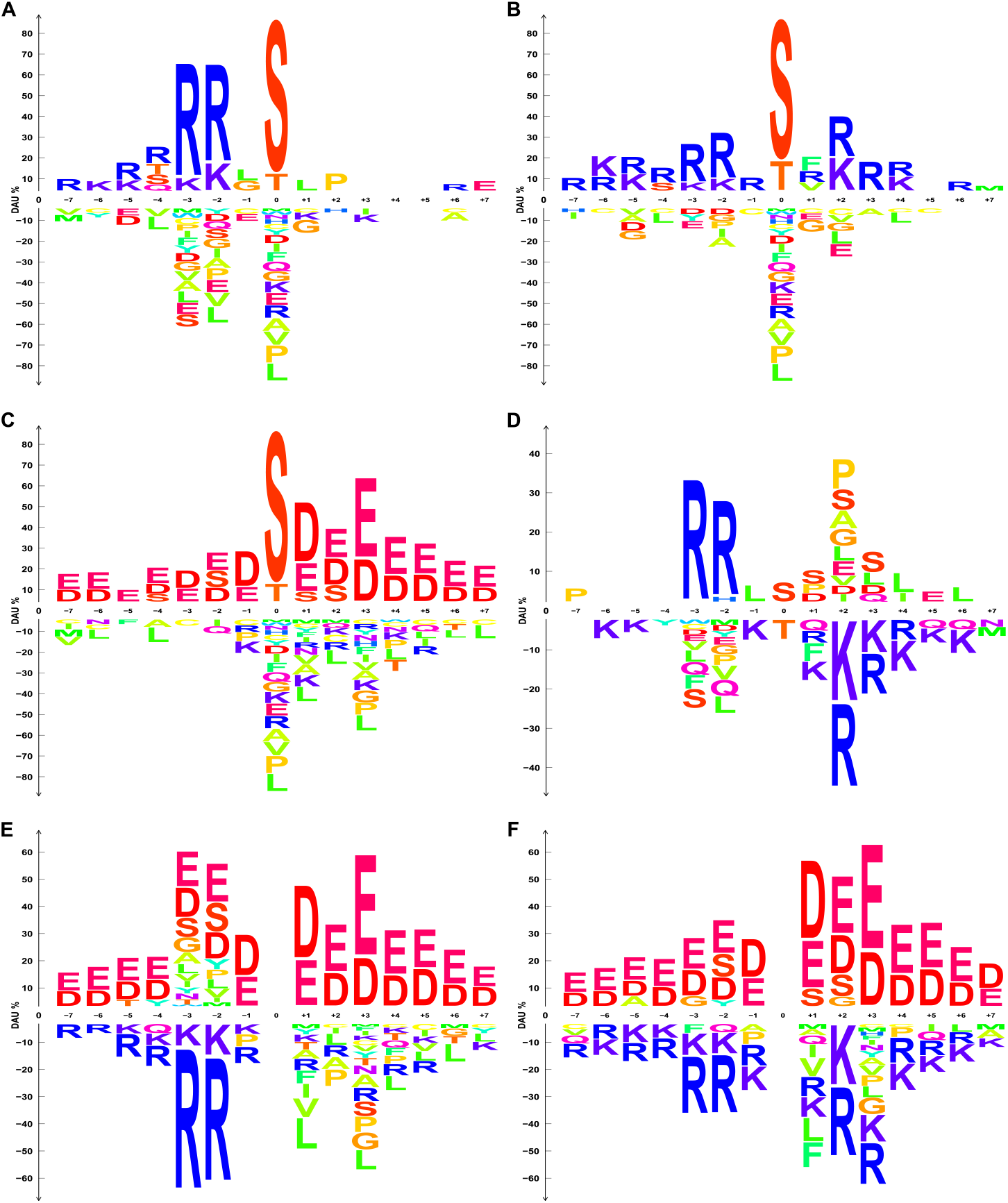
Sequence logos showing the substrate sequence specificities of human Ser/Thr kinases. (**A-C**) Logos showing substrate phosphorylation motifs of human PKACA, PKCA, and CK2A1, respectively. A background model was built from randomly sampled subsequences of 15 AA residues from the UniProt human reference proteome for *Z*-test with a significance level of 0.05. DAU tests were performed to identify AA residue preferences in human substrate phosphorylation motifs of human PKACA, PKCA, and CK2A1 separately. Over- and under-represented AA residues with significant DAU at each pattern position are shown as sequence logos. Individual AA residues are colored differently. (**D-F**) Logos showing differential substrate AA residue preferences of PKACA over PKCA, CK2A1 over PKACA, and CK2A1 over PKCA, respectively. Subsequences of 15 AA residues (−7 to +7), centered on the substrate phosphorylation sites of two kinases to compare, were used as the input set and the background set, respectively. Fisher’s exact test was performed with a significance level of 0.05.

Next, we compared the substrate preferences of the three human STKs using all known substrates curated by the PhosphoSitePlus database, with the subsequences of each other’s substrates as background sets. The differential phosphorylation site analyses further indicated that the substrate motif of PKACA is much more similar to that of PKCA than to that of CK2A1 (**Figure 2**). PKCA prefers basic residues at the *C*-terminal of the phosphorylation sites more than PKACA does, while PKACA prefers basic residues at positions −2 and −3 more than PKAC does, which is evident in **Figure 2D.** Degenerate logos displaying differential usage of AA residues grouped by charge under physiological conditions further highlight the differential substrate preferences and similarities of the three STKs in terms of charge of residues flanking phosphorylation sites (**Supplementary Figure S2**). The frequencies of Thr and Ser residues at the phosphorylation sites (position 0) of PKCA and PKACA substrates are significantly different (**Figure 2D**). This example demonstrated that dagLogo enables the discovery of charge preferences around the phosphorylation sites of human STKs with an AA grouping and the differential AA preferences among different kinases.

### Case study 3: dagLogo confirms substrate specificity of the yeast *N*-terminal acetyltransferase

*N*-terminal acetylation of proteins by *N*-terminal acetyltransferases (Nat) is evolutionarily conserved from bacteria to humans [71–74]. More than 80% of human proteins and more than 50% of yeast proteins are *N*-terminal acetylated [75, 76]. Different *N*-terminal acetyltransferases (NatA to NatG) have different substrate preferences depending on the *N*-terminal AA residue context, especially the first and second residues, of mature proteins [71, 73]. About 57% and 84% of *N*-terminal acetylation of yeast and human proteins are mediated by NatA [75], which preferentially acetylates proteins with Ser-, Ala-, Thr-, Val- or Gly-V-termini [71, 75].

We reanalyzed the yeast NatA substrate specificity using dagLogo with the first 25 residues of yeast NatA substrate as the input set and the first 25 *N*-terminal residues of yeast proteins excluding the NatA substrates as the background set. Fisher’s exact test was performed to determine the significantly differential usage of AAs in the yeast NatA substrate motif. Our analysis demonstrated that the yeast NatA preferentially acetylates proteins with Ser, followed by Ala, as the first residue at the mature *N*-termini, which is consistent with previous studies [7, 75, 77, 78] (**Figure 3A**). However, the original publications [71, 75] also reported that proteins with mature *N*-termini starting with Thr, Val, or Gly residue are preferred by NatA, although no statistical test was performed. In contrast, our analysis does not support such findings (**Figure 3A**), which is consistent with a reanalysis of the data with a proper background model using iceLogo [7]. Besides the first residues, the residues at positions 2 to 25 also seem to be involved in determining substrate specificity. Glutamine (Q) residues are preferred at multiple positions of the *N*-terminal sequences. By grouping AAs based on charge, we also found that yeast NatA strongly prefers uncharged AA residues at position 1 and slightly prefers negatively charged residues at positions 2, 3, 6, 10, 11, 17, 20, and 23, but prefers positively charge residues at position 12 (**Figure 3B**). Positively charged residues are disliked at positions 1 to 6, and uncharged residues are disfavored at positions 11 and 12 (**Figure 3B**). When grouping AAs based on size, we observed that yeast NatA strongly prefers tiny residues as the starting AA. In addition, tiny to medium size AAs are favored, and large to gigantic residues are disliked at multiple positions (**Figure 3C**).

**Figure 3.**
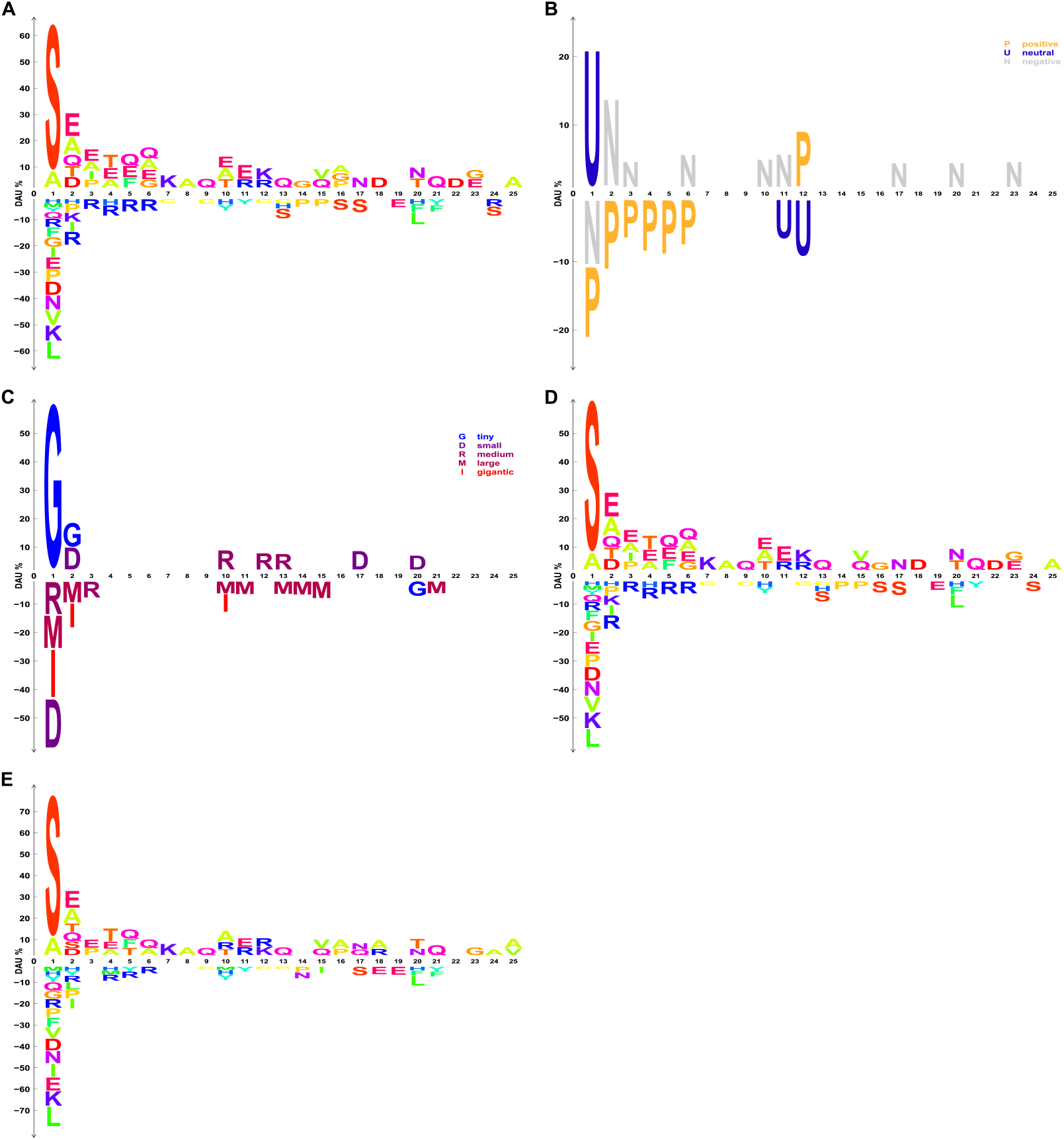
Sequence logos showing the *N*-terminal sequence pattern of the yeast NatA substrates. (**A–C**) A background model was built from the first 25 *N*-terminal residues of known yeast, non-NatA substrates for Fisher’s exact test with a significance level of 0.05. (**A**) Over- and under-represented AA residues with significant DAU at each pattern position are shown as sequence logos. Individual AA residues are colored differently. (**B** and **C**) AA residues in both the input and background sets are grouped, collapsed, and represented by single letters based on their charge status under physiological conditions and sizes, respectively. Positively charged AAs = {H, K, R}, neutral AAs = {A, C, F, G, I, L, M, N, P, Q, S, T, V, W, Y}, and negatively charged AAs = {D, E}. Tiny = {G, S, A}, small = {D, N, C, E, H}, medium = {R, Q, K, T}, large = {M, P, Y, F}, and gigantic = {I, L, V, W}. Fisher’s exact test was performed with a significance level of 0.05. Over- and under-represented AA residue groups with significant DAU at each pattern position are shown as sequence logos. AA residue groups are colored according to their charge status (**B**) and sizes (**C**). (**D** and **E**) Background models for Fisher’s exact test with a significance level of 0.05 were built from the first 25 *N*-terminal residues of all yeast proteins and from randomly sampled subsequences of 25 AA residues from the yeast reference proteome, respectively.

Next, we investigated how different background models affect the results of differential AA usage analysis of the yeast NatA substrate motifs. Background models were built using a background set of the first 25 *N*-terminal residues of all yeast proteins and a background set of randomly sampled subsequences of 25 AA residues from the yeast reference proteome separately. **Figures 3D** and **3E** show differential AA usage for NatA substrate motifs given the two different background models. It is evident that the identities and percentages of differential usage of some AAs at multiple positions are different when different background models were adopted (**Figures 3D** and **3E** versus **Figure 3A**). This result underscores the importance of choosing the correct background model based on the study objectives and the advantage of dagLogo.

### Comparison of statistical test methods for differential AA usage analyses

We compared how different statistical test methods, *Z*-test and Fisher’s exact test, affect differential AA usage analysis results. For this purpose, we analyzed the human GRB substrate data [46] and the yeast NatA substrate data [75] to represent both ends of the proteome size spectrum with the human proteome (74,823 proteins) as a large and the yeast proteome (6,049 proteins) as a small proteome. Although Fisher’s exact test runs slower than z-test for the larger proteome, results based on both methods are largely similar to each other regardless of proteome size (**Supplementary Figure S3**). However, the identities and percentage of differential usage of some AAs are slightly different (**Supplementary Figure S3**).

## Discussion

To facilitate the discovery of conserved patterns from increasingly accumulated proteomic data, we developed the dagLogo package. Our package helps to identify and visualize differentially used AA residues or AA residue groups in given protein sequences, improving and extending the sophisticated logo generator iceLogo. First, dagLogo corrects the way of calculating standard deviation of position-specific AA frequencies implemented by iceLogo. In iceLogo, standard deviation of 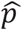 is not correctly computed; it should be 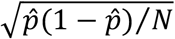 instead of 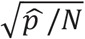, where 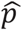 is the sample estimate of the proportion of a given AA residue at a given pattern position, and *N* is the total number of sequences in a background set [7]. Second, dagLogo implements both *Z*-test and Fisher’s exact test, whereas iceLogo only implements *Z*-test. Fisher’s exact test computes the exact p-value, while Z-test uses a normal distribution as an approximation of a binomial distribution, which becomes problematic when the background sets or input sets are small [79]. When the background sets are large, such as the whole human proteome, both tests produce very similar results, though the Z-test-based method is faster than the Fisher’s exact test-based one. However, our analysis also shows that the identities and percentages of differential usage of some AAs can be different (**Supplementary Figure S3**).

Unlike most logo generators, our tool allows multiple different types of inputs. These flexible options can easily eliminate users’ efforts to prepare aligned subsequences. Peptide sequences identified by proteomic assays can be directly used as input for pattern discovery. As far as we know, only RNALogo can perform alignment if the input is not pre-aligned [9]. More importantly, twelve background models can be easily generated for *Z*-test or Fisher’s exact test using dagLogo. It is very important to choose an appropriate background model based on the biological questions and experimental protocols. Using the yeast NatA substrate data, we demonstrated the importance of choosing the most appropriate background model (**Figure 3A** and **3D**). Of note, to build the most relevant background model, users may need to focus only on proteins expressed under a given condition, in a specific tissue or cell type, or even in a subcellular compartment.

Additionally, in logos generated by all existing tools, the occurrence of individual residue at a given position is displayed either without considering statistical significance or without allowing degenerate representation based on residue similarity. dagLogo provides more than ten grouping schemes to lump together AA residues of similar physicochemical properties—such as charge, hydrophobicity, and size—into single-letter symbols at given positions. In addition, the function *addScheme* can be used for adding more customized grouping schemes. AA groupings reduce the AA alphabet for testing and visualization. Tests of differential usage of grouped AAs can increase the chances of discovering and visualizing subtle differences that might not be detectable or obvious at the individual AA level (**Figures 1, 2** and **3**).

## Conclusion

Re-analyses of three datasets using dagLogo demonstrated that dagLogo with appropriate background model can recapitulate previous findings about the substrate motifs of human GRB, three human Ser/Thr kinases, and yeast NatA by allowing flexible input formats, context-specific background models, and AA groupings. Most importantly, analysis with dagLogo at the AA group level revealed important physicochemical property of substrate preferences that would have been hidden with the analysis at the individual AA level alone.

## Materials and methods

### Implementation

dagLogo is implemented as an open source R/Bioconductor [80] package. Several S4 classes are implemented to represent different datatypes: Class *dagPeptides* for peptide sequences, Class *Proteome* for proteomes, Class *dagBackground* for background models, and Class *testDAUresults* for statistical test results of differential usage of AA residues/groups. Functions implemented in the dagLogo package are described in **Supplementary Table S2** and the package vignette.

### Workflow of dagLogo analyses

A flowchart for a typical dagLogo-based analysis is shown in **Figure 4**. A dagLogo analysis can start with creating a *Proteome* object, which stores protein identifiers (IDs), protein sequences, species information, and the source of sequences. One can prepare the *Proteome* object with the *prepareProtome* function by providing a fasta file containing the proteome sequences of the species. Alternatively, the user can specify the species’ scientific name or NCBI taxonomy ID with the same function, which will automatically download the species-specific proteome data from the UniProt database.

**Figure 4.**
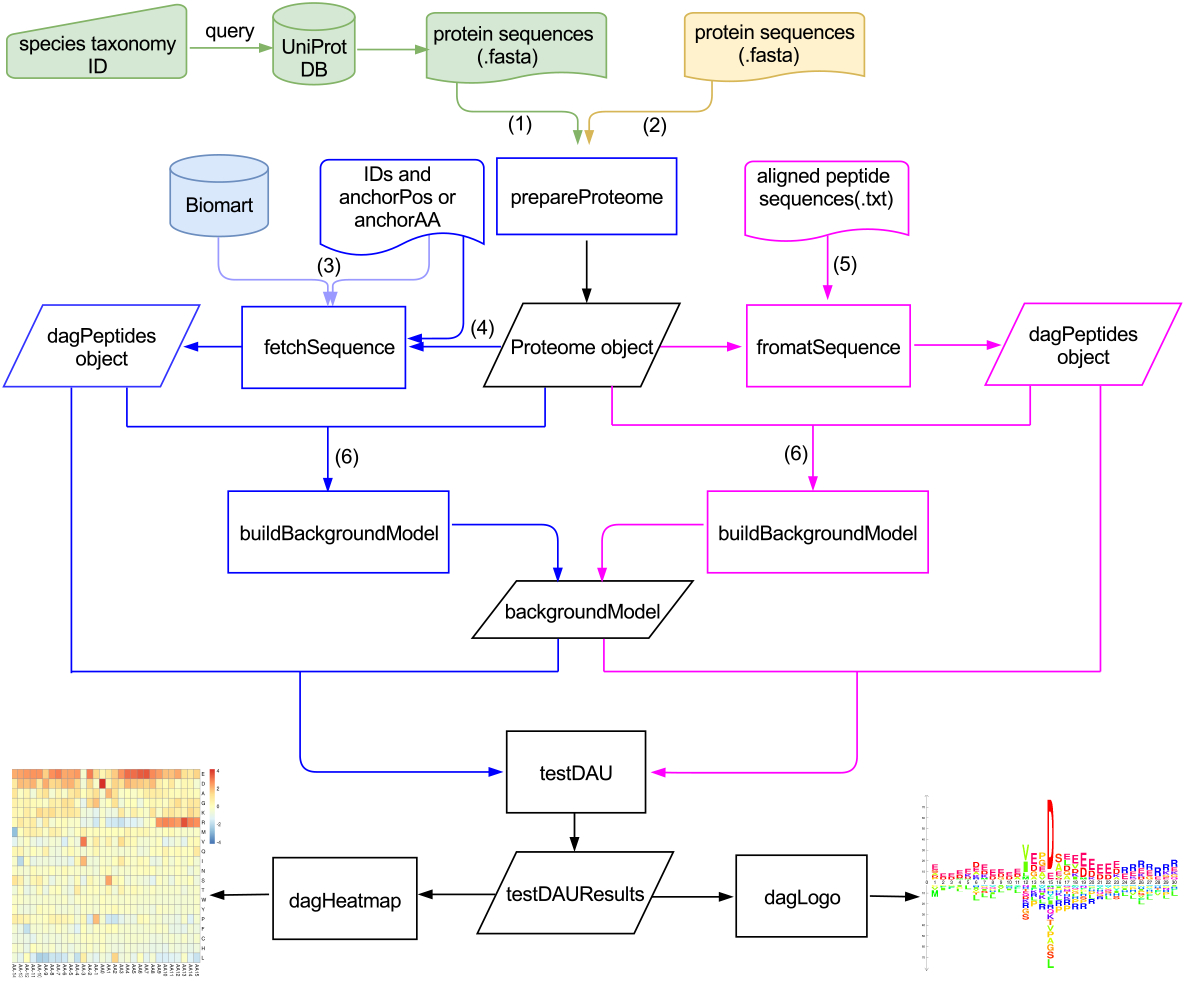
dagLogo workflow. When whole proteome sequences in the fasta format are not available for a species, the NCBI taxonomy ID or scientific name of the species is needed to query the UniProt database to obtain the desired fasta file (1) and to build a *Proteome* object using the *prepareProteomeByFTP* or *prepareProteomeByUniprotws* function. These two functions are wrapped into the *prepareProteome* function. Otherwise, an existing fasta file (2) is directly used to build a *Proteome* object with the *prepareProteome* function. A *dagPeptides* object for an input set can be constructed in three ways: i) Entrez gene IDs or UniProt SwissProt protein IDs, as well as anchoring AAs (such as “K” for Lys) or anchoring AA position information of the input set, can be used to obtain sequences from the species-specific Biomart database for creating *a dagPeptides* object using the *fetchSequence* function. An anchoring AA within a peptide sequence is represented by a lowercase single-letter symbol or an uppercase single-letter symbol followed by an asterisk, e.g., the anchoring AA Lys in “ASTRSkSSTD” or “ASTRSK*SSTD”. Meanwhile, an anchoring AA position is represented as a string consisting of the anchoring AA followed by the AA’s position, such as “K123” for the 123th residue — Lys (3); ii) With the same input information as the previous method, a *dagPeptides* object can be built from the species-specific *Proteome* object rather than the Biomart database using the *fetchSequence* function (4); iii) If the peptide sequences of the input set have already been aligned, the *dagPeptides* object can be built from the species-specific Proteome object using the *formatSequence* function. Twelve different background models for Fisher’s exact test or *Z*-test can be built from the *Proteome* object using the *buildBackground* function with different parameter settings (6). A differential AA or AA group usage (DAU) analysis for the input set (the dagPetides object) can be performed using the *testDAU* function, with the appropriate background model represented by a *backgroundModel* object). Heatmaps and logos generated by the functions *dagHeatmap* and *dagLogo,* respectively, can be used to visualize the results of the DAU analysis.

Next, a *dagPeptides* object representing a formatted input set can be created in several ways. For pre-aligned input subsequences, one can use the *formatSequence* function with the specified *proteome* object, as well as upstream and downstream offsets relative to the anchoring positions of the input subsequences. If pre-aligned input subsequences are unavailable, the *dagPeptides* object can be created by using the *fetchSequence* function in four different ways by providing IDs of protein sequences from which input subsequences originate, the options to access full-length protein sequences from an Ensembl Biomart database or a *proteome* object, the anchoring AA(s) and the anchoring AA’s positions. Prior to fetching sequences, input data can be cleaned up with the *cleanPeptides* function, which can remove peptides without anchoring AAs or duplicated peptides, and can generate an individual entry for each anchoring AA if an original entry contains multiple anchoring AAs.

Like iceLogo, dagLogo requires a background model, which is derived from background subsequences of the same length as those in the input set, to properly calculate the probability of the number of occurrences of given AAs at each position in the input set. Choosing an appropriate background model is therefore of great importance, and the background set should suit the actual technical and biological contexts. Twelve background models can be generated for *Z*-test or Fisher’s exact test (see below) using the *buildBackgroundModel* function by specifying i) the proteome-level background-generating space as the whole proteome (“WholeProteome”), proteins corresponding to the input set (“InputSet”), or the whole proteome excluding those proteins in “InputSet” (“NonInputSet”); ii) the protein-level background-generating space as *N*-termini only, *C*-termini only, or anywhere of full-length proteins; iii) anchoring AAs or not when the protein-level background-generating space is not restricted to protein termini. A proper background model should be considered to meet experimental and analytical needs [7]. Uniport, Biomart, and/or anchored AAs are leveraged to fetch the peptide sequences, which have characteristics or constraints similar to those of the experimental data, to build a specific background model. For example, if an experiment only focuses on the mitochondrial proteome, then an appropriate background model would be built from the entire or a subset of the mitochondrial proteome that is anchored to the AA(s) of interest. If the experiment is to identify peptides with acetylated lysine, then an appropriate background model would be derived from all peptide fragments containing anchoring lysine.

Once the type of background model to build is decided, a Fisher’s exact test or *Z*-test approximation can be applied to identify significantly enriched/depleted AAs or AA groups. For Fisher’s exact test, a single background set with all available background subsequences will be generated. If both the input set’s size and the number of subsequences for the background set are large enough, *Z*-test approximation may be preferred, as it can speed up the tests of differential AA or AA group usage (DAU). To do so, bootstrapping is used to generate multiple background sets that conform to the background-generating rules, with each background set consisting of the same number of equal-length sequences as those in the input set. The averaged frequency 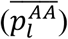 of a given residue *AA* at position *l* and the corresponding standard error 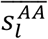 are calculated as follows:

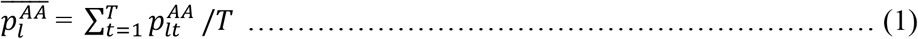

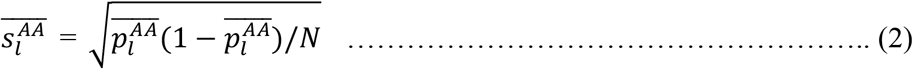

where 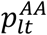 is the frequency of residue *AA* at position *l* for subsample *t,* with *I* ∈ [1, *L*] and *t*∈ [1, *T*]. *L* is the length of the pattern, *T* is the total number of subsamples of background sets, and *N* is the number of subsequences in each subsample. Notably, the number of sequences in each background set and the total number of background sets can affect the accuracy of the estimates for AA frequencies and their standard errors. A normal distribution 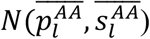 is used to approximate the distribution of the proportion of a given residue at each position.

Next, given the background model built as above, the significance of the proportion of an AA at a given pattern position in the experimental set is tested by using the *testDAU* function. Performing Fisher’s exact test is straightforward. To perform a two-sided *Z*-test, we assume the frequency of a residue at a given pattern position of the input set approximately follows the normal distribution 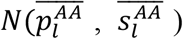, which is inferred from the background sets for Z-test. A Z-score is calculated as follows:

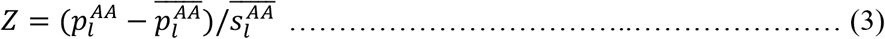

where 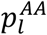 is the frequency of a given *AA* at given position *l* in the input set. The test result is an object of Class *testDAUresults*. The test results can be visualized as a heatmap using the *dagHeatmap* function, which leverages the *pheatmap* package. Significantly over- and under-represented residues can be visualized as sequence logos using the dagLogo function, which is built on the grid graphics system in R.

Importantly, the package also provides an array of grouping schemes to represent AA residues of similar properties with degenerate, single-letter symbols. AAs can be grouped according to the indices of their isoelectric point (pI) [81], polarity [82], hydrophobicity [83, 84], bulkiness [81], volume [85], and consensus similarity [86] as individual AAs or substitutability within protein contexts [87–89]. In addition, dagLogo allows users to specify customized grouping and coloring schemes through the *addScheme* function. Users can refer to the AAindex database [40] or other resources [26, 90] to create their own grouping schemes. With grouped AAs represented by reduced AA alphabets, dagLogo can improve the detection power of identifying significant degenerate residues and can generate degenerate sequence logos.

### Datasets for case studies

We demonstrated the applications of our dagLogo package with three datasets as described below.

#### Dataset 1: Substrates of the human GRB

We downloaded the Supplementary Table S1 by Van Damme *et al.* [46], and made the following modifications: i) four obsolete UniProtKB-TrEMBL identifiers (IDs) were changed to the current UniProt Swiss-Prot IDs of proteins with identical sequences, as determined by BLAST analyses; ii) IDs of three proteins with accession numbers starting with “IPI” were changed to their current UniProt Swiss-Prot IDs; and iii) six substrates without current Swiss-Prot records (O55106, O95365, P78559, Q00839, Q06210 and Q149F1) were removed. These manipulations resulted in 416 human protein substrates of the human GRB for the case study. The processed data are included as **Supplementary Table S3.** Equal-length subsequences (P15-P15’) centered on the human GRB cutting sites—the peptide bonds between P1-P1’—were prepared using the *fetchSequence* function in the dagLogo package. Herein P15 to P1 are residues at positions 15 to 1 *N*-terminal to the cutting sites, while P1’ to P15’ are residues at position 1 to 15 *C*-terminal to the cutting sites. The letter X was used to pad the peptide sequences when the fetched sequence’s length was shorter than 30. These 416 subsequences were used as the input set to build a dagPeptides object. The background set consisted of the 300 subsets of 416 subsequences of 30 AA residues randomly sampled from the UniProt human reference proteome (https://www.uniprot.org/proteomes/UP000005640).

#### Dataset 2: Substrates of human Ser/Thr protein kinases

We downloaded phosphorylated peptide sequences from all known kinase substrates from the PhosphoSitePlus database (https://www.phosphosite.org/) (file name: “Kinase_Substrate_Dataset.gz”). Then, we extracted the UniProt IDs, peptide sequences of 15 AAs (phosphorylation site ± 7 AAs), and absolute positions of phosphorylation sites of 867, 646, and 651 substrates of the human kinases PKACA, PKCA and CK2A1 from 11 species, including the human. Of these sequences, 469, 385, and 465 are human substrates of the human kinases PKACA, PKCA, and CK2A1, respectively. The data are included as **Supplementary Table S4.** To determine the human substrate sequence specificities of each human kinase, the same background model was built using randomly sampled subsequences of 15 AAs from the UniProt human reference proteome (https://www.uniprot.org/proteomes/UP000005640). To compare the substrate sequence preference of one kinase with that of another, multi-species substrate subsequences of 15 AAs (phosphorylation site ± 7 AAs) of one kinase were used as the input set, and the multi-species substrate subsequences of 15 AAs (phosphorylation site ± 7 AAs) of another kinase were used as the background set. *Dataset 3: Substrates of the yeast N-terminal acetyltransferase A (NatA)*

The latest baker’s yeast reference proteome was downloaded from the UniProt database (https://www.uniprot.org/proteomes/UP000002311). The *N*-terminal Met residue was removed from each protein sequence in the entire proteome for subsequent analysis for two reasons: i) the first Met residue is cleaved co-translationally *in vivo* if the next residue is glycine, alanine, serine, threonine, cysteine, proline, or valine [91]; ii) all eukaryotic NatA does not acetylate a peptide or protein with a Met *N*-terminus [71, 75, 77, 78]. The data used for mining the substrate sequence specificity of yeast NatA was extracted from the Supplementary Table S3 with the following modifications [75]. Obsolete IDs of six substrates (P04451, P26781, P26782, P35271, P40213, and P53030) were updated to the current UniProt Swiss-Prot IDs. Misinformation of the mature *N*-terminal 5-AA peptide sequences (column name: “mature_N_term_P5”) of 11 substrates (Q12074, P02829, P00360, P38720, P04076, P15019, P10591, Q03048, P53880, P38088, and P35187) were corrected based on the yeast reference proteome. With these manipulations, the final data consisted of information about 285 substrates of the yeast NatA. Peptide sequences consisting of the first 25-AA of the *N*-termini were retrieved from the yeast proteome using the *fetchSequence* function in the dagLogo package. The data are included as **Supplementary Table S5.** Different background sets (see **Results**) were adopted depending on the analysis purposes.

## Authors’ contributions

LJZ conceived of and coordinated the study. JO, HL, and LJZ designed and developed the software. HL analyzed the data. HL and LJZ interpretated data and wrote the first draft. NKN and UA provided feedback on amino acid groupings and initial data for testing. AS contributed functions to the software. All authors read, edited, and approved the final manuscript.

## Acknowledgements

This work was supported by internal funding from the Department of Molecular, Cell and Cancer Biology in the University of Massachusetts Medical School. We thank Serena Han for editorial assistance.

## Supplementary material

**Figure S1.**
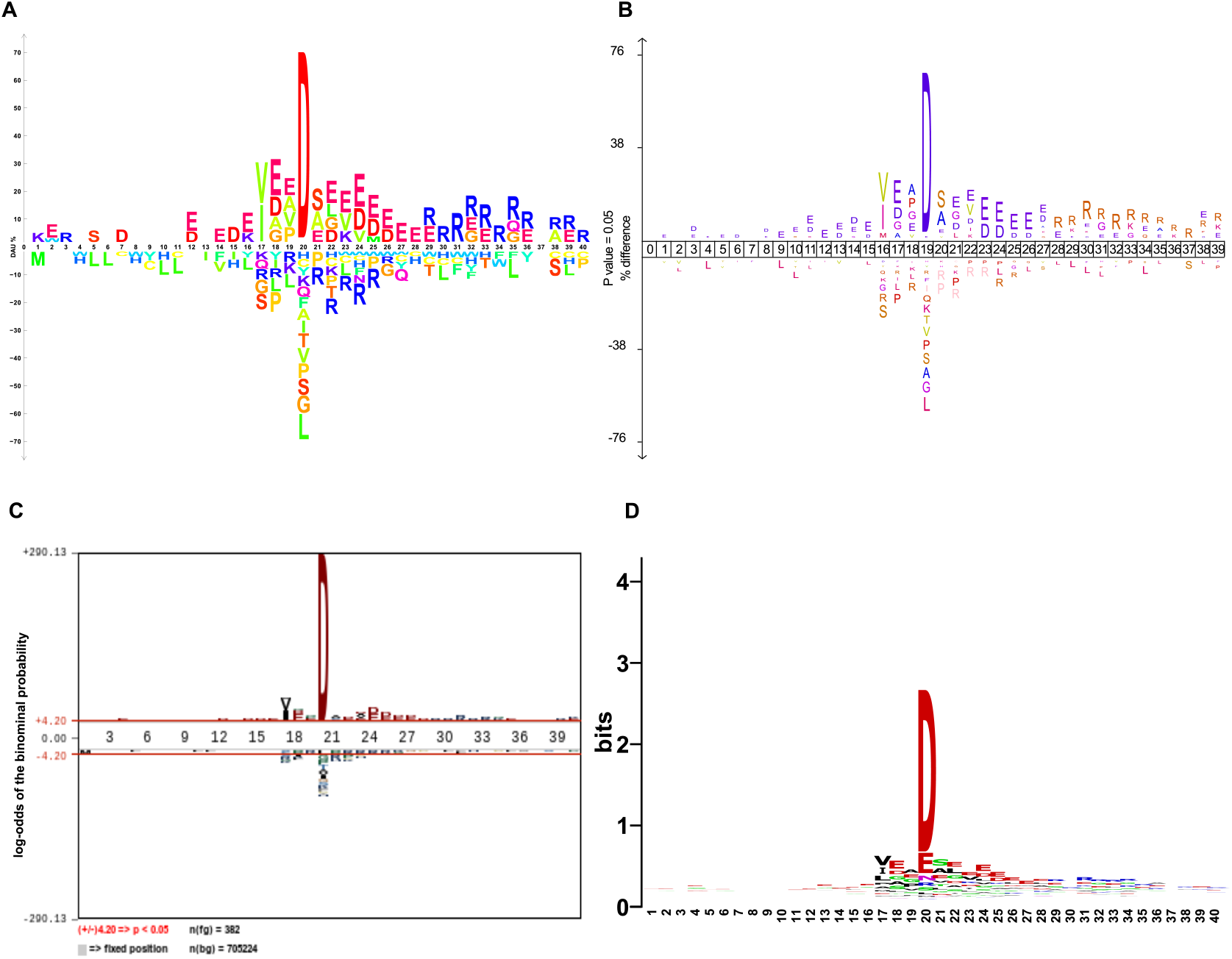
Sequences logos generated by different tools showing sequence determinants of human GRB substrate specificity. The same set of 416 aligned, equal-length subsequences of 30 residues centered on the cleavage sites was used as the input set. The same set of randomly sampled subsequences of 30 AA residues from the human reference proteome was used as the background for iceLogo, pLogo, and dagLogo. A uniform global background of equal probability (0.05) of each of the 20 natural AAs was used for WebLogo. (**A**) dagLogo, (**B**) iceLogo, (**C**) pLogo, and (**D**) WebLogo.

**Figure S2.**
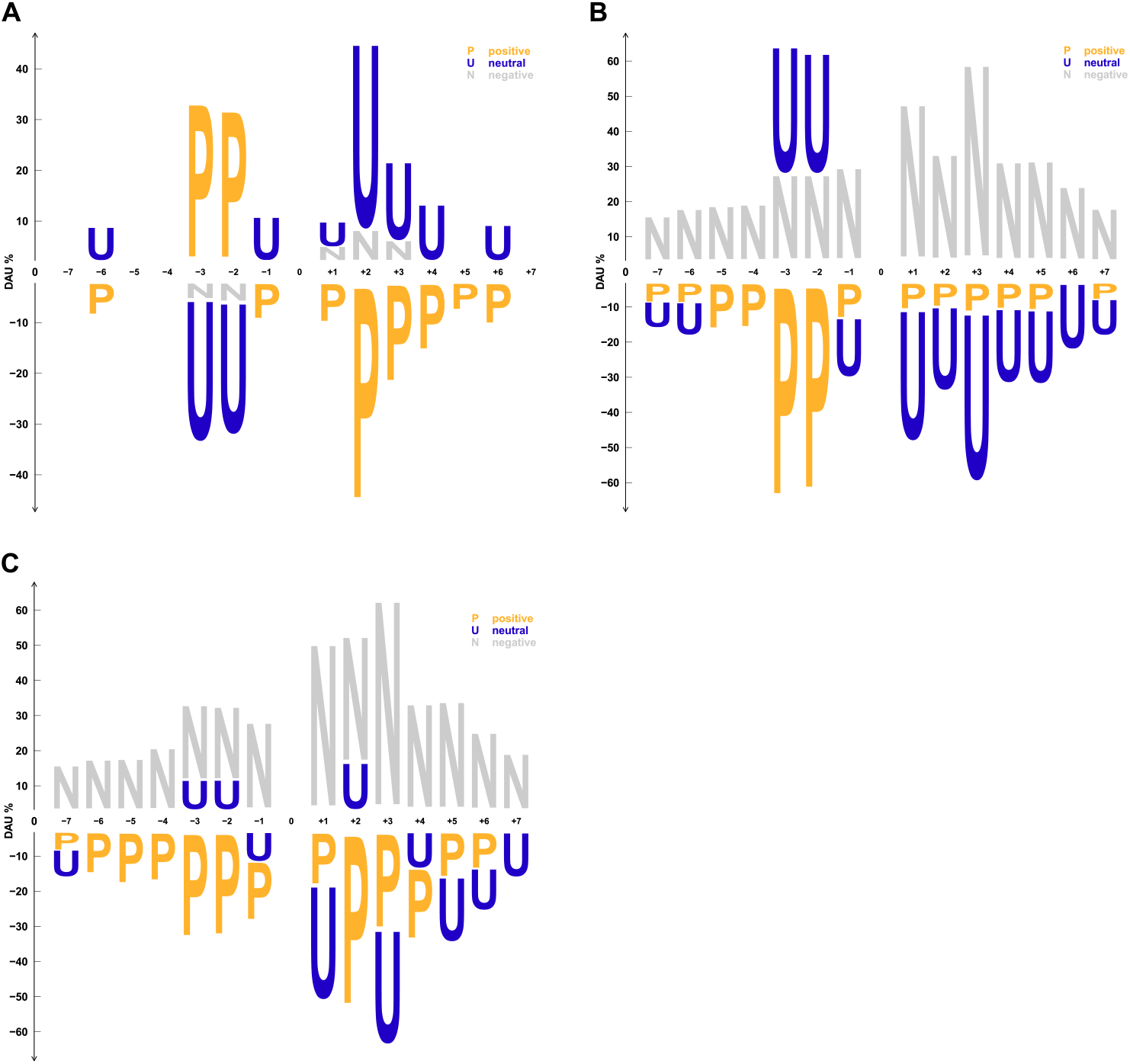
Degenerate dagLogos showing differential AA group usage in substrate motifs of human STKs. Subsequences of 15 AA residues (−7 to +7) centered on the substrate phosphorylation sites of two kinases were used as the input set and the background set, respectively. AA residues were grouped by their charge status under physiological conditions, and the differential usage of AA groups was tested by using the function *testDAU* to perform Fisher’s exact test with a significance level of 0.05. Positively charged AAs = {H, K, R}, neutral AAs = {A, C, F, G, I, L, M, N, P, Q, S, T, V, W, Y}, and negatively charged AAs = {D, E}. (**A-C**) Logos showing differential substrate AA residue group preferences of PKACA over PKCA, CK2A1 over PKACA, and CK2A1 over PKCA, respectively.

**Figure S3.**
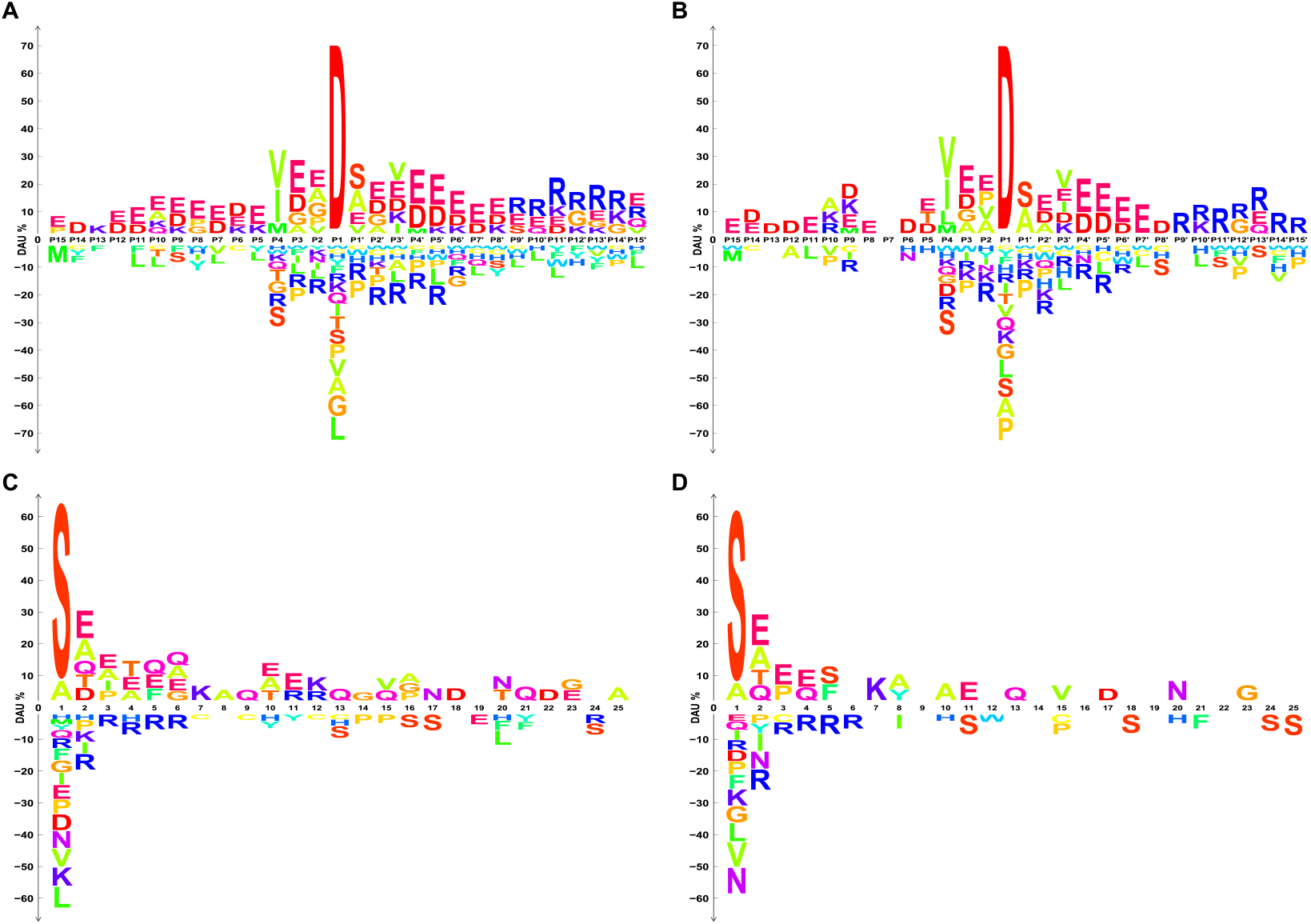
dagLogos resulting from different statistical test methods. Fisher’s exact test (**A** and **C**) and *Z*-test (**B** and **D**) were performed to identify substrate sequence preferences of human GRB and yeast NatA, with a significance level of 0.05. For human GRB substrate specificity analyses, the same set of 416, equal-length subsequences of 30 residues centered on the cleavage sites was used as the input set, while background models for Fisher’s exact test and *Z*-test were built from all subsequences and from randomly sampled subsequences of 30 AA residues from the UniProt human reference proteome, respectively. For yeast NatA substrate specificity analyses, the same set of subsequences of 25 residues from the *N*-termini of the 285 NatA substrates was used as the input set, while background models for Fisher’s exact test and *Z*-test were built from all subsequences and from randomly sampled subsequences of 25 AA residues from the *N*-termini of the yeast proteins not including the 285 NatA substrates, respectively. The significance level was set at 0.05 for all tests.

**Table S1 A comprehensive comparison of existing logo generators**

**Table S2 Description of main functions implemented in the dagLogo package**

**Table S3 Information about substrates of human GRB for case study 1**

**Table S4 Information about substrates of human Ser/Thr kinases for case study 2**

**Table S5 Information about substrates of yeast NatA for case study 3**

## References

[1] Gorodkin J, Heyer LJ, Brunak S, Storomo GD. Displaying the in formation contents of structural RNA alignments: the structure logos. Bioinformatics 1997;13:583–6.

[2] Crooks GE, Hon G, Chandonia JM, Brenner SE. WebLogo: a sequence logo generator. Genome Res 2004;14:1188–90.

[3] Schneider TD, Stephens RM. Sequence logos: a new way to display consensus sequences. Nucleic Acids Res 1990;18:6097–100.

[4] Dey KK. A Brief History of Sequence Logos. Biostatistics and Biometrics Open Access Journal 2018;6:102–5.

[5] Wheeler TJ, Clements J, Finn RD. Skylign: a tool for creating informative, interactive logos representing sequence alignments and profile hidden Markov models. BMC Bioinformatics 2014;15:7–.

[6] O’Shea JP, Chou MF, Quader SA, Ryan JK, Church GM, Schwartz D. pLogo: a probabilistic approach to visualizing sequence motifs. Nat Methods 2013;10:1211–2.

[7] Colaert N, Helsens K, Martens L, Vandekerckhove J, Gevaert K. Improved visualization of protein consensus sequences by iceLogo. Nat Methods 2009;6:786–7.

[8] Workman CT, Yin Y, Corcoran DL, Ideker T, Stormo GD, Benos PV. enoLOGOS: a versatile web tool for energy normalized sequence logos. Nucleic Acids Res 2005;33:389.

[9] Chang TH, Horng JT, Huang HD. RNALogo: a new approach to display structural RNA alignment. Nucleic Acids Res 2008;36:91.

[10] Li W, Yang B, Liang S, Wang Y, Whiteley C, Cao Y, et al. BLogo: a tool for visualization of bias in biological sequences. Bioinformatics 2008;24:2254–5.

[11] Ye Z, Ma T, Kalmbach MT, Dasari S, Kocher JA, Wang L. CircularLogo: a lightweight web application to visualize intra-motif dependencies. BMC Bioinformatics 2017;18:269–2.

[12] Thomsen MC, Nielsen M. Seq2Logo: a method for construction and visualization of amino acid binding motifs and sequence profiles including sequence weighting, pseudo counts and two-sided representation of amino acid enrichment and depletion. Nucleic Acids Res 2012;40:281.

[13] Vacic V, Iakoucheva LM, Radivojac P. Two Sample Logo: a graphical representation of the differences between two sets of sequence alignments. Bioinformatics 2006;22:1536–7.

[14] Schreiber M, Brown C. Compensation for nucleotide bias in a genome by representation as a discrete channel with noise. Bioinformatics 2002;18:507–12.

[15] Foat BC, Morozov AV, Bussemaker HJ. Statistical mechanical modeling of genome-wide transcription factor occupancy data by MatrixREDUCE. Bioinformatics (Oxford, England) 2006;22:141.

[16] Shannon CE. A mathematical theory of communication. Bell Syst. Tech. J. 1948;27:379–423.

[17] Kullback S, Leibler RA. On information and sufficiency. Ann. Math. Statist. 1951;22:79–86.

[18] Schneider TD, Stormo GD, Gold L, Ehrenfeucht A. Information content of binding sites on nucleotide sequences. J Mol Biol 1986;188:415–31.

[19] Dey KK, Xie D, Stephens M. A new sequence logo plot to highlight enrichment and depletion. BMC Bioinformatics 2018;19:473.

[20] Hasan S, Schreiber M. Recovering motifs from biased genomes: application of signal correction. Nucleic Acids Res 2006;34:5124–32.

[21] Maddelein D, Colaert N, Buchanan I, Hulstaert N, Gevaert K, Martens L. The iceLogo web server and SOAP service for determining protein consensus sequences. Nucleic Acids Res 2015;43:543.

[22] Wu X, Bartel DP. kpLogo: positional k-mer analysis reveals hidden specificity in biological sequences. Nucleic Acids Res 2017.

[23] Saethang T, Hodge K, Yang CR, Zhao Y, Kimkong I, Knepper MA, et al. PTM-Logo: a program for generation of sequence logos based on position-specific background amino-acid probabilities. Bioinformatics 2019;35:5313–4.

[24] Chan HS. Folding alphabets. Nat Struct Biol 1999;6:994–6.

[25] Wang J, Wang W. A computational approach to simplifying the protein folding alphabet. Nat Struct Biol 1999;6:1033–8.

[26] Bacardit J, Stout M, Hirst JD, Valencia A, Smith RE, Krasnogor N. Automated alphabet reduction for protein datasets. BMC Bioinformatics 2009;10:6.

[27] Huang JT, Wang T, Huang SR, Li X. Reduced alphabet for protein folding prediction. Proteins 2015;83:631–9.

[28] Zheng L, Liu D, Yang W, Yang L, Zuo Y. RaacLogo: a new sequence logo generator by using reduced amino acid clusters. Brief. Bioinform. 2020.

[29] Pratt H, Weng Z. LogoJS: a Javascript package for creating sequence logos and embedding them in web applications. Bioinformatics 2020.

[30] Ye Y, Choi JH, Tang H. RAPSearch: a fast protein similarity search tool for short reads. BMC Bioinformatics 2011;12:159.

[31] Li T, Fan K, Wang J, Wang W. Reduction of protein sequence complexity by residue grouping. Protein Engineering, Design and Selection 2003;16:323–30.

[32] Melo F, Marti-Renom MA. Accuracy of sequence alignment and fold assessment using reduced amino acid alphabets. Proteins 2006;63:986–95.

[33] Biro JC. Amino acid size, charge, hydropathy indices and matrices for protein structure analysis. Theoretical Biology and Medical Modelling 2006;3:15.

[34] Li J, Wang W. Grouping of amino acids and recognition of protein structurally conserved regions by reduced alphabets of amino acids. Sci China C Life Sci 2007;50:392–402.

[35] Bacardit J, Stout M, Hirst JD, Sastry K, Llorà X, Krasnogor N (2007), ‘Automated alphabet reduction method with evolutionary algorithms for protein structure prediction’, Proceedings of the 9th annual conference on Genetic and evolutionary computation, Association for Computing Machinery, London, England, pp.346–53.

[36] Peterson EL, Kondev J, Theriot JA, Phillips R. Reduced amino acid alphabets exhibit an improved sensitivity and selectivity in fold assignment. Bioinformatics (Oxford, England) 2009;25:1356–62.

[37] Zheng L, Huang S, Mu N, Zhang H, Zhang J, Chang Y, et al. RAACBook: a web server of reduced amino acid alphabet for sequence-dependent inference by using Chou’s five-step rule. Database (Oxford) 2019;2019:baz131.

[38] Hu X, Friedberg I. SwiftOrtho: a fast, memory-efficient, multiple genome orthology classifier. Gigascience 2019;8.

[39] Amano M, Hamaguchi T, Shohag MH, Kozawa K, Kato K, Zhang X, et al. Kinase-interacting substrate screening is a novel method to identify kinase substrates. The Journal of cell biology 2015;209:895–912.

[40] Kawashima S, Pokarowski P, Pokarowska M, Kolinski A, Katayama T, Kanehisa M. AAindex: amino acid index database, progress report 2008. Nucleic Acids Res 2008;36:D202–5.

[41] Betts MJ, Russell RB. Amino acid properties and consequences of substitutions. In: Gray M. R. B. a. I. C. (ed) Bioinformatics for Geneticists. John Wiley & Sons, Ltd, 2003, 289–316.

[42] Masson D, Tschopp J. A family of serine esterases in lytic granules of cytolytic T lymphocytes. Cell 1987;49:679–85.

[43] Trapani JA. Granzymes: a family of lymphocyte granule serine proteases. Genome Biol 2001;2:reviews3014.1.

[44] Poe M, Blake JT, Boulton DA, Gammon M, Sigal NH, Wu JK, et al. Human cytotoxic lymphocyte granzyme B. Its purification from granules and the characterization of substrate and inhibitor specificity. J Biol Chem 1991;266:98–103.

[45] Harris JL, Peterson EP, Hudig D, Thornberry NA, Craik CS. Definition and redesign of the extended substrate specificity of granzyme B. J Biol Chem 1998;273:27364–73.

[46] Van Damme P, Maurer-Stroh S, Plasman K, Van Durme J, Colaert N, Timmerman E, et al. Analysis of protein processing by N-terminal proteomics reveals novel species-specific substrate determinants of granzyme B orthologs. Mol. Cell. Proteom. 2009;8:258–72.

[47] Waugh SM, Harris JL, Fletterick R, Craik CS. The structure of the pro-apoptotic protease granzyme B reveals the molecular determinants of its specificity. Nat Struct Biol 2000;7:762–5.

[48] Ruggles SW, Fletterick RJ, Craik CS. Characterization of structural determinants of granzyme B reveals potent mediators of extended substrate specificity. J Biol Chem 2004;279:30751–9.

[49] Estébanez-Perpiña E, Fuentes-Prior P, Belorgey D, Braun M, Kiefersauer R, Maskos K, et al. Crystal structure of the caspase activator human granzyme B, a proteinase highly specific for an Asp-P1 residue. Biol Chem 2000;381:1203–14.

[50] Cohen P. The origins of protein phosphorylation. Nat Cell Biol 2002;4:E127–E30.

[51] Pawson T, Scott JD. Protein phosphorylation in signaling-50 years and counting. Trends Biochem Sci 2005;30:286–90.

[52] Hunter T. Protein kinases and phosphatases: the yin and yang of protein phosphorylation and signaling. Cell 1995;80:225–36.

[53] Ubersax JA, Ferrell Jr JE. Mechanisms of specificity in protein phosphorylation. Nat Rev Mol Cell Biol 2007;8:530–41.

[54] Zhu G, Liu Y, Shaw S. Protein kinase specificity. A strategic collaboration between kinase peptide specificity and substrate recruitment. Cell Cycle 2005;4:52–6.

[55] Amanchy R, Periaswamy B, Mathivanan S, Reddy R, Tattikota SG, Pandey A. A curated compendium of phosphorylation motifs. Nat Biotechnol 2007;25:285–6.

[56] Rust HL, Thompson PR. Kinase consensus sequences: a breeding ground for crosstalk. ACS Chem Biol 2011;6:881–92.

[57] Kemp BE, Pearson RB. Protein kinase recognition sequence motifs. Trends Biochem Sci 1990;15:342–6.

[58] Cheng F, Jia P, Wang Q, Zhao Z. Quantitative network mapping of the human kinome interactome reveals new clues for rational kinase inhibitor discovery and individualized cancer therapy. Oncotarget 2014;5:3697–710.

[59] Mok J, Kim PM, Lam HYK, Piccirillo S, Zhou X, Jeschke GR, et al. Deciphering Protein Kinase Specificity Through Large-Scale Analysis of Yeast Phosphorylation Site Motifs. Science Signaling 2010;3:ra12–ra.

[60] Capra M, Nuciforo PG, Confalonieri S, Quarto M, Bianchi M, Nebuloni M, et al. Frequent alterations in the expression of serine/threonine kinases in human cancers. Cancer Res 2006;66:8147–54.

[61] Ardito F, Giuliani M, Perrone D, Troiano G, Lo Muzio L. The crucial role of protein phosphorylation in cell signaling and its use as targeted therapy (Review). Int J Mol Med 2017;40:271–80.

[62] Olsen JV, Blagoev B, Gnad F, Macek B, Kumar C, Mortensen P, et al. Global, in iivo, and site-specific phosphorylation dynamics in signaling networks. Cell 2006;127:635–48.

[63] Giansanti P, Stokes MP, Silva JC, Scholten A, Heck AJ. Interrogating cAMP-dependent kinase signaling in Jurkat T cells via a protein kinase A targeted immune-precipitation phosphoproteomics approach. Mol Cell Proteomics 2013;12:3350–9.

[64] Hennrich ML, Marino F, Groenewold V, Kops GJ, Mohammed S, Heck AJ. Universal quantitative kinase assay based on diagonal SCX chromatography and stable isotope dimethyl labeling provides high-definition kinase consensus motifs for PKA and human Mps1. J Proteome Res 2013;12:2214–24.

[65] Nishikawa K, Toker A, Johannes FJ, Songyang Z, Cantley LC. Determination of the specific substrate sequence motifs of protein kinase C isozymes. J Biol Chem 1997;272:952–60.

[66] Meggio F, Marin O, Pinna LA. Substrate specificity of protein kinase CK2. Cell Mol Biol Res 1994;40:401–9.

[67] Gnad F, Young A, Zhou W, Lyle K, Ong CC, Stokes MP, et al. Systems-wide analysis of K-Ras, Cdc42, and PAK4 signaling by quantitative phosphoproteomics. Mol Cell Proteomics 2013;12:2070–80.

[68] Rusin SF, Adamo ME, Kettenbach AN. Identification of candidate casein kinase 2 substrates in mitosis by quantitative phosphoproteomics. Front. Cell Dev. Biol. 2017;5:97.

[69] Bian Y, Ye M, Wang C, Cheng K, Song C, Dong M, et al. Global screening of CK2 kinase substrates by an integrated phosphoproteomics workflow. Sci Rep 2013;3:3460.

[70] St-Denis N, Gabriel M, Turowec JP, Gloor GB, Li SS, Gingras AC, et al. Systematic investigation of hierarchical phosphorylation by protein kinase CK2. J Proteomics 2015;118:49–62.

[71] Aksnes H, Drazic A, Marie M, Arnesen T. First things first: vital protein marks by N-terminal acetyltransferases. Trends Biochem Sci 2016;41:746–60.

[72] Ouidir T, Jarnier F, Cosette P, Jouenne T, Hardouin J. Characterization of N-terminal protein modifications in Pseudomonas aeruginosa PA14. J Proteomics 2015;114:214–25.

[73] Linster E, Wirtz M. N-terminal acetylation: an essential protein modification emerges as an important regulator of stress responses. J Exp Bot 2018;69:4555–68.

[74] Soppa J. Protein acetylation in archaea, bacteria, and eukaryotes. Archaea 2010;2010:820681.

[75] Arnesen T, Van Damme P, Polevoda B, Helsens K, Evjenth R, Colaert N, et al. Proteomics analyses reveal the evolutionary conservation and divergence of N-terminal acetyltransferases from yeast and humans. Proc Natl Acad Sci U S A 2009;106:8157–62.

[76] Van Damme P, Hole K, Pimenta-Marques A, Helsens K, Vandekerckhove J, Martinho RG, et al. NatF contributes to an evolutionary shift in protein N-terminal acetylation and is important for normal chromosome segregation. PLoS Genet 2011;7:e1002169–e.

[77] Polevoda B, Sherman F. N-terminal acetyltransferases and sequence requirements for N-terminal acetylation of eukaryotic proteins. J Mol Biol 2003;325:595–622.

[78] Polevoda B, Norbeck J, Takakura H, Blomberg A, Sherman F. Identification and specificities of N-terminal acetyltransferases from *Saccharomyces cerevisiae*. EMBO J 1999;18:6155–68.

[79] McDonald JH. Handbook of Biological Statistics Baltimore, Maryland.: Sparky House Publishing, 2014.

[80] Gentleman RC, Carey VJ, Bates DM, Bolstad B, Dettling M, Dudoit S, et al. Bioconductor: open software development for computational biology and bioinformatics. Genome Biol 2004;5:R80.

[81] Zimmerman JM, Eliezer N, Simha R. The characterization of amino acid sequences in proteins by statistical methods. J.Theor. Biol. 1968;21:170–201.

[82] Grantham R. Amino acid difference formula to help explain protein evolution. Science 1974;185:862–4.

[83] Kyte J, Doolittle RF. A simple method for displaying the hydropathic character of a protein. J Mol Biol 1982;157:105–32.

[84] Hopp TP, Woods KR. Prediction of protein antigenic determinants from amino acid sequences. Proc Natl Acad Sci U S A 1981;78:3824–8.

[85] Bigelow CC. On the average hydrophobicity of proteins and the relation between it and protein structure. J Theor Biol 1967;16:187–211.

[86] Stephenson JD, Freeland SJ. Unearthing the root of amino acid similarity. J Mol Evol 2013;77:159–69.

[87] Dayhoff M, Schwartz R, Orcutt B. A model of evolutionary change in proteins. In: Dayhoff M. (ed) Atlas of Protein Sequence and Structure. Washington, D. C.: National Biomedical Research Foundation, 1978, 345–52.

[88] Mirny LA, Shakhnovich EI. Universally conserved positions in protein folds: reading evolutionary signals about stability, folding kinetics and function. J Mol Biol 1999;291:177–96.

[89] Maiorov VN, Crippen GM. Contact potential that recognizes the correct folding of globular proteins. J Mol Biol 1992;227:876–88.

[90] Kosiol C, Goldman N, Buttimore NH. A new criterion and method for amino acid classification. J Theor Biol 2004;228:97–106.

[91] Wingfield PT. N-terminal methionine processing. Current protocols in protein science 2017;88:6.14.1–6.14.3.

